# The transcription factor *Si*ROCX controls a monocot-specific CAZyme program for host cell wall remodeling during symbiotic root colonization

**DOI:** 10.64898/2026.03.05.709870

**Authors:** Laura Armbruster, Mathias Brands, Leo Baumgart, Frederic Dreesmann, Lisa Leson, Vicente Ramírez, Yu Zhang, Taim Nassr, Gregor Langen, Markus Pauly, Alga Zuccaro

## Abstract

Intracellular accommodation of beneficial fungi requires controlled remodeling of host cell walls while avoiding activation of plant immune responses. During colonization of monocot roots, the endophytic fungus *Serendipita indica* induces a suite of carbohydrate-active enzymes (CAZymes) targeting xylan and cellulose. Weighted gene co-expression network analysis identified the basidiomycete-specific transcription factor *Si*ROCX as a key regulator of this monocot-adapted program. DNA affinity purification sequencing (DAP-seq) defined the *Si*ROCX binding motif and revealed a CAZyme-enriched target regulon. Inducible *Si*ROCX overexpression selectively activated motif-containing CAZyme genes and markedly enhanced xylanase and cellulase activities on xylan and native barley root cell walls. Premature activation of this program triggered expression of a plant immune marker despite comparable fungal biomass, indicating that precise temporal control of CAZyme deployment is required to maintain symbiotic compatibility. Together, these findings identify *Si*ROCX as a central regulator of a monocot-adapted cell wall–degrading secretome and reveal a basidiomycete-specific regulatory module that coordinates host cell wall remodeling with immune-compatible symbiotic colonization.

**Significance:** Beneficial fungi colonize living plant cells, requiring host cell wall remodeling while avoiding immune activation. However, the key transcriptional regulators coordinating this process in basidiomycetes during symbiotic root colonization remain unknown. We identify *Si*ROCX, a conserved basidiomycete-specific transcription factor, as a master activator of a monocot-adapted xylan/cellulose degradation program in the root endophyte *Serendipita indica*. By integrating *in planta* co-expression networks, DAP-seq, secretome proteomics, and enzymatic assays, we show that *Si*ROCX overexpression markedly enhances secretion and activity of xylan- and cellulose-degrading enzymes, boosting sugar release from monocot cell walls. These findings reveal a basidiomycete-specific regulatory module for immune-compatible host cell wall remodeling, providing a framework to engineer fungal CAZyme programs for crop symbiosis and biomass conversion.

## Introduction

Plants interact with a wide variety of soil-borne microorganisms that they recruit from the bulk soil into the rhizosphere (Bulgarelli et al., 2012; Lareen et al., 2016; Yang et al., 2024). The root cell wall serves as both a protective barrier against pathogens and a dynamic interface for beneficial microbes (Underwood, 2012). To overcome this barrier and establish themselves within host tissues, root-colonizing fungi secrete a diverse suite of carbohydrate-active enzymes (CAZymes) that degrade host cell wall polysaccharides. Collectively, these enzymes facilitate fungal penetration, nutrient acquisition, and the processing of immunogenic carbohydrates of plant or microbial origin that could otherwise elicit host immune responses (Kubicek et al., 2014; Hage and Rosso, 2021).

CAZymes are grouped into several functional classes, including glycoside hydrolases (GHs), glycosyl transferases (GTs), polysaccharide lyases (PLs), carbohydrate esterases (CEs), and enzymes with auxiliary activities (AAs). Together, these enzymes break down the major polysaccharide components of plant biomass, cellulose, hemicelluloses, and pectins (Campbell et al., 1997; Henrissat and Davies, 1997; Coutinho et al., 2003; Boraston et al., 2004; Lombard et al., 2010; Jiménez et al., 2025). Cellulose, the most abundant plant polysaccharide, is a linear polymer of 1,4-β-linked glucose units arranged into crystalline microfibrils (French, 2017; Marinho, 2025). Its hydrolysis requires the concerted action of AAs, which loosen the crystalline structure, and GHs, primarily exo-glucanases, endo-glucanases, and β-glucosidases, which subsequently cleave the glucan chains (Henriksson et al., 2000; Cantarel et al., 2009; French, 2017; Andlar et al., 2018; Marinho, 2025). Hemicelluloses are branched polysaccharides composed of diverse pentoses and hexoses. Their abundance and composition differ substantially between dicot and monocot cell walls. While monocot walls are rich in xylans and mixed-linkage glucans, dicot walls primarily contain xyloglucans, along with smaller amounts of glucomannans and mannans (Fry, 1989; Carpita and Gibeaut, 1993; Vogel, 2008; O’Neill and York, 2018). As a result of this structural diversity, hemicellulose breakdown requires a broad suite of CEs, AAs and GHs, including xylanases, mannanases, arabinases and glucanases (Hage and Rosso, 2021). Pectin is a heteropolysaccharide composed mainly of galacturonic acid and rhamnose, with branched side chains of arabinose, galactose or xylose. Compared to dicot cell walls, grass cell walls contain relatively little pectin (Mohnen, 2008; Szatanik-Kloc et al., 2017). Pectin depolymerization is facilitated by GHs, in particular polygalacturonases and rhamnogalacturonases, together with PLs and CEs (Voragen et al., 2009; Li et al., 2024).

The composition and size of fungal CAZyme repertoires reflect their host range and colonization strategies (King et al., 2011; Zhao et al., 2013; Hage and Rosso, 2021). Biotrophic fungi, which derive nutrients from living host cells, typically harbour reduced CAZyme sets to limit host damage and avoid triggering defense responses. In contrast, necrotrophic and hemibiotrophic fungi actively kill host cells to meet their nutritional demands and therefore generally encode larger and more diverse CAZyme repertoires (Zhao et al., 2013). Similarly, generalist fungi such as *Botrytis cinerea* tend to harbour enlarged CAZyme repertoires that provide them with the flexibility to break down both monocot and dicot cell walls efficiently (Nagel et al., 2021).

In addition to variations in CAZyme repertoire size and composition, functional diversity also emerges from differences in the regulation of CAZyme expression. Comparative analyses of closely related endophytic and parasitic fungi revealed that, despite harbouring largely similar CAZyme repertoires, parasitic species exhibit earlier, stronger and more diversified CAZyme induction *in planta*, suggesting they employ a more aggressive strategy for extracting carbon from host cell walls than their endophytic counterparts (Hacquard et al., 2016; Muñoz-Barrios et al., 2020; Moonjely and Trail, 2025). Even within a single fungal species, transitions between lifestyles are accompanied by pronounced shifts in CAZyme expression, reflecting tight transcriptional control. The beneficial root endophyte *Serendipita indica* (*Si*) for instance has an extended CAZyme repertoire resembling that of soil decomposers and wood-decaying fungi, reflecting its saprotrophic ancestry (Zuccaro et al., 2011; Gong et al., 2022). Consistent with this, *Si* follows a biphasic colonization strategy, beginning with a biotrophic phase and transitioning to a saprotrophic phase at later stages. Although the latter involves localized host cell death, the damage remains tightly constrained and does not result in root necrosis, allowing the interaction to remain biotrophic at the tissue level (Lahrmann et al., 2015; Weiß et al., 2016). The transition from the biotrophic to the saprotrophic phase is accompanied by a pronounced induction of CAZyme expression, highlighting the dynamic regulation of cell wall–modifying enzymes throughout the colonization process (Lahrmann et al., 2015; Eichfeld et al., 2024; Brands et al., 2025).

Apart from colonization stage, fungi also fine-tune CAZyme expression to match the polysaccharide composition and architecture of the respective host cell wall (Zhao et al., 2013; Hage and Rosso, 2021; Brands et al., 2025). Transcriptome profiling of *Rhizoctonia solani* during infection of rice, maize and soybean showed that CAZyme expression patterns markedly differ during the infection of the different hosts, demonstrating an adaptation of the pathogen to the different cell wall compositions (Xia et al., 2017). Similarly, *Si* as a generalist beneficial fungus that colonizes monocots and dicots alike adjusts its CAZyme expression profile in a host-dependent manner (Eichfeld et al., 2024).

The principal regulatory mechanism governing CAZyme expression is carbon catabolite repression (CCR). CCR suppresses CAZyme transcription in the presence of easily metabolizable carbon sources such as glucose (Gancedo, 1998; Adnan et al., 2017). In addition, several conserved transcriptional activators of CAZyme genes have been identified in ascomycetes, among them the Zn(II)_2_Cys_2_ transcription factors AraR (Battaglia et al., 2011), ClrA/CLR-1 (Raulo et al., 2016), ClrB/CLR-2/ManR (Ogawa et al., 2012; Raulo et al., 2016; Kun et al., 2023), RhaR (Gruben et al., 2014) and XlnR (van Peij et al., 1998; Raulo et al., 2016). Notably, none of these regulators have orthologs in basidiomycetes, suggesting that the regulation of plant biomass degradation emerged after the evolutionary split between ascomycetes and basidiomycetes (Rytioja et al., 2014; Todd et al., 2014).

The fact that basidiomycetes show synchronized upregulation of CAZymes during growth on plant biomass (Martinez et al., 2009; Vanden Wymelenberg et al., 2009; MacDonald et al., 2011; Brands et al., 2025) suggests they rely on regulatory systems functionally analogous to those found in ascomycetes. Recent work revealed that *S. indica* employs a coordinated enzymatic strategy to degrade acetylated xylan in monocot roots, involving sequential activity of the endo-1,4-β-xylanase *Si*GH11 and acetyl-xylan esterase *Si*AXE (Brands et al., 2025). Notably, this enzymatic program is activated during later stages of colonization, coinciding with the onset of localized host cell death. However, the transcriptional regulators coordinating this CAZyme program during symbiotic colonization remain unknown. Here, we investigate the regulation of CAZyme induction in the basidiomycete *S. indica* during barley colonization. Combining time-resolved transcriptomics, DAP-seq, and functional analyses, we identify *Si*ROCX, a basidiomycete-specific transcription factor that coordinates xylan- and cellulose-degrading CAZymes during late-stage monocot colonization. Our results show that *Si*ROCX drives the production of a host cell wall–degrading secretome and that its precise temporal activation is critical for enabling host cell wall remodeling while maintaining symbiotic compatibility.

## Results

### Discovery of the *Si*ROCX CAZyme regulon during monocot colonization

Building on time-resolved RNA-seq of *S. indica* colonizing *Hordeum vulgare* (*Hv*) and *Brachypodium distachyon* (*Bd*) (Eichfeld et al 2024), weighted gene co-expression network analysis (WGCNA) (Brands et al., 2025) identified two monocot-induced modules (M4, M7) enriched for secreted CAZymes (Fig. 1A).

**Fig. 1.**
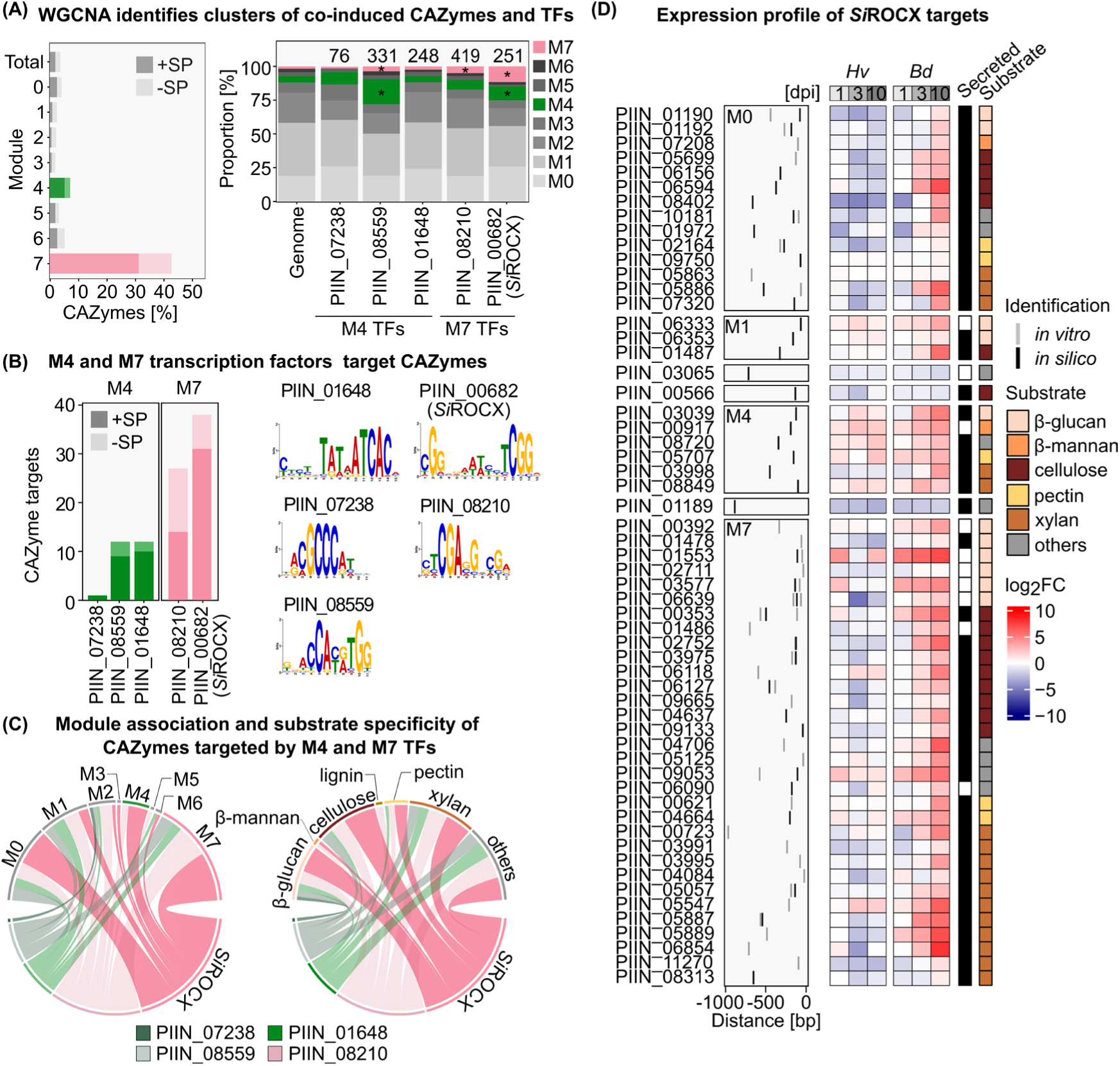
*Si*ROCX regulates the expression of host cell wall-degrading carbohydrate-active enzymes (CAZymes). **A)** Weighted gene co-expression network analysis (WGCNA) of time-resolved RNA-seq data from *Serendipita indica* (*Si*) during colonization of *Hordeum vulgare* (*Hv*) and *Brachypodium distachyon* (*Bd*) identified seven modules of co-expressed genes, including two monocot-induced modules (M4 and M7) enriched for secreted CAZymes. The distribution of DAP-seq–identified transcription factor (TF) target genes across modules is shown for selected candidate TFs. Modules enriched (>2-fold relative to the genome) are marked by an asterisk. **B)** DAP-seq identified CAZyme targets and enriched binding motifs for selected M4- and M7-associated TFs. The number of CAZyme targets per TF is shown on the left, and enriched binding motifs are visualized as sequence logos representing nucleotide enrichment at each position. Among the profiled TFs, *Si*ROCX (PIIN_00682) bound the largest number and highest proportion of CAZyme promoters. **C)** Chord diagrams illustrate module association and predicted substrate specificity of CAZymes targeted by M4- and M7-associated TFs (substrate annotations adapted from Brands et al., 2025). **D)** Genomic localization of *Si*ROCX binding sites relative to the predicted start codon, predicted substrate specificity, and expression profiles of *Si*ROCX target CAZymes during host colonization. Expression profiles during colonization of *Hv* and *Bd* are shown as log2 fold change values along with prediction of secretion and predicted substrate. In addition to direct DAP-seq targets (*in vitro*), further candidate targets were identified by integrating motif scanning (FIMO; MEME Suite v5.5.5; Bailey et al., 2009) with regulatory network inference (GENIE3; Huynh-Thu et al., 2010) (*in silico*).

We hypothesized that coordinated induction of CAZymes within these modules is driven by transcriptional master regulators. Using high module membership scores (>0.75), we identified 11 transcription factors (TFs) as candidate regulators of M4 and 3 TFs as candidate regulators of M7 (Supplementary Table S1). Nine candidates were subjected to DNA-affinity purification sequencing (DAP-seq), in which *in vitro*–synthesized Halo-tagged TFs were incubated with sheared *S. indica* genomic DNA, and TF-bound fragments were recovered and sequenced. Four TFs did not pass quality control (≤100,000 filtered aligned fragments; FRIP ≤0.01) and were excluded from further analysis. For the remaining five TFs, genes with DAP-seq peaks within 1 kb upstream of the predicted start codon were defined as high-confidence targets, yielding 76 (PIIN_07238) to 419 (PIIN_08210) targets per TF (Fig. 1A). Target enrichment analyses supported the WGCNA-based regulator assignments. Although M4 genes represent 4.8% of the genome, they comprised 19% of targets of the M4-associated TF PIIN_08559 (62/331; ∼4-fold enrichment). Likewise, M7 genes (1.7% of the genome) were enriched among targets of PIIN_08210 (21/419; 5%; ∼3-fold enrichment) and PIIN_00682 (29/251; 11.5%; ∼7-fold enrichment). Together, these enrichments indicate that WGCNA reliably grouped candidate TFs with their downstream targets.

Among the TFs with high-confidence binding motifs (Fig. 1B, Supplementary Table S2), PIIN_00682 targeted the largest number of CAZymes both in absolute and relative terms (38 CAZymes among 251 total targets; 15%; Supplementary Table S3). Functional annotation of PIIN_00682 targets showed a strong bias toward CAZymes involved in degradation of plant cell wall components, including β-glucans, cellulose, and xylan (Fig. 1C). We therefore refer to PIIN_00682 as REGULATOR OF CELLULASES AND XYLANOLYTIC ENZYMES (*SiROCX*). To expand the *SiROCX* regulon beyond high-confidence DAP-seq targets, we searched promoter regions (1 kb upstream of the start codon) for the *SiROCX* binding motif using FIMO (MEME suite) (Bailey et al., 2009; Grant et al., 2011). In parallel, we inferred regulatory interactions from the colonization RNA-seq dataset using GENIE3 (Huynh-Thu et al., 2010). Genes were classified as additional *SiROCX* targets if they harboured a binding motif in the 1 kb upstream region of their predicted start codon and were linked to *SiROCX* in the GENIE3 network (Supplementary Table S4). This *in silico* integration identified 20 additional putative CAZyme targets, the majority of which were strongly induced during monocot colonization (Fig. 1D).

Next, to place *Si*ROCX in an evolutionary context, we reconstructed a maximum-likelihood phylogeny of fungal Zn₂Cys₆ transcription factors from Ascomycota and Basidiomycota using IQ-TREE2 (Supplementary Fig. 1). The phylogeny included 114 fungal-specific Zn₂Cys₆ transcription factors reported by Marian et al. (2022), defined by the presence of both a Zn₂Cys₆ DNA-binding domain (Pfam PF00172) and a fungal-specific transcription factor domain (Pfam PF04082), together with eight additional sequences closely related to *Si*ROCX and representative Zn₂Cys₆ family members. The cellulase regulator Roc1 from the saprotrophic wood-decaying fungus *Schizophyllum commune*, conserved within Agaricomycetes, served as a reference to delineate a Roc1/ROCX-like lineage.

To distinguish Roc1/ROCX-like candidates within this family, we applied a custom profile HMM built from Agaricomycete Roc1/ROCX sequences; sequences with domain scores >500 were classified as Roc1/ROCX(Marian et al., 2022). As shown, *Si*ROCX (778 aa) groups within the Basidiomycota-specific clade (grey background) and clusters tightly with *S. commune* Roc1 (bootstrap 92%), distinct from ascomycete CAZyme regulators (e.g., XlnR, ClrA/CLR-1, ClrB/CLR-2), which fall below the HMM threshold. Combined with Roc1’s confinement to Agaricomycetes and its association with lignocellulose degradation, these findings indicate that Roc1/ROCX transcription factors form a Basidiomycota-specific regulatory lineage distinct from canonical ascomycete CAZyme regulators. This lineage appears to have been adapted to diverse ecological contexts, ranging from saprotrophic wood decay to biotrophic plant colonization.

### *Si*ROCX activates a monocot-adapted xylan/cellulose-degrading secretome

To validate candidate *Si*ROCX targets *in vivo*, we overexpressed *SiROCX* under the glucose-inducible *Si*FGB1 promoter (Wawra et al., 2016), confirmed growth of the resulting transformants under cell wall stress conditions, and quantified transcript levels of *SiROCX* and six CAZyme genes selected as representative SiROCX targets in axenic culture by qRT-PCR (Fig. 2A and Supplementary Fig. 2). Three CAZymes were selected based on experimental identification as *Si*ROCX targets via DAP-seq (PIIN_00353, PIIN_06639, PIIN_09053), and three were inferred as targets based on motif and network analysis (FIMO/GENIE3; PIIN_03995, PIIN_04706, PIIN_06118). These genes represent diverse CAZyme families (AA9, CE4, GH1, GH5, GH6, GH62) and predicted substrate classes. In the absence of the host, CAZyme expression was low in the wild-type strain, whereas *Si*ROCX overexpression significantly upregulated all six target genes in at least one transformant relative to wild type (WT). Induction magnitude correlated with *SiROCX* expression across three independent overexpression strains (*Si*ROCX-OX #1–#3), which showed approximately 10-, 20-, and 45-fold increases in *Si*ROCX transcript abundance (R² > 0.7).

**Fig. 2.**
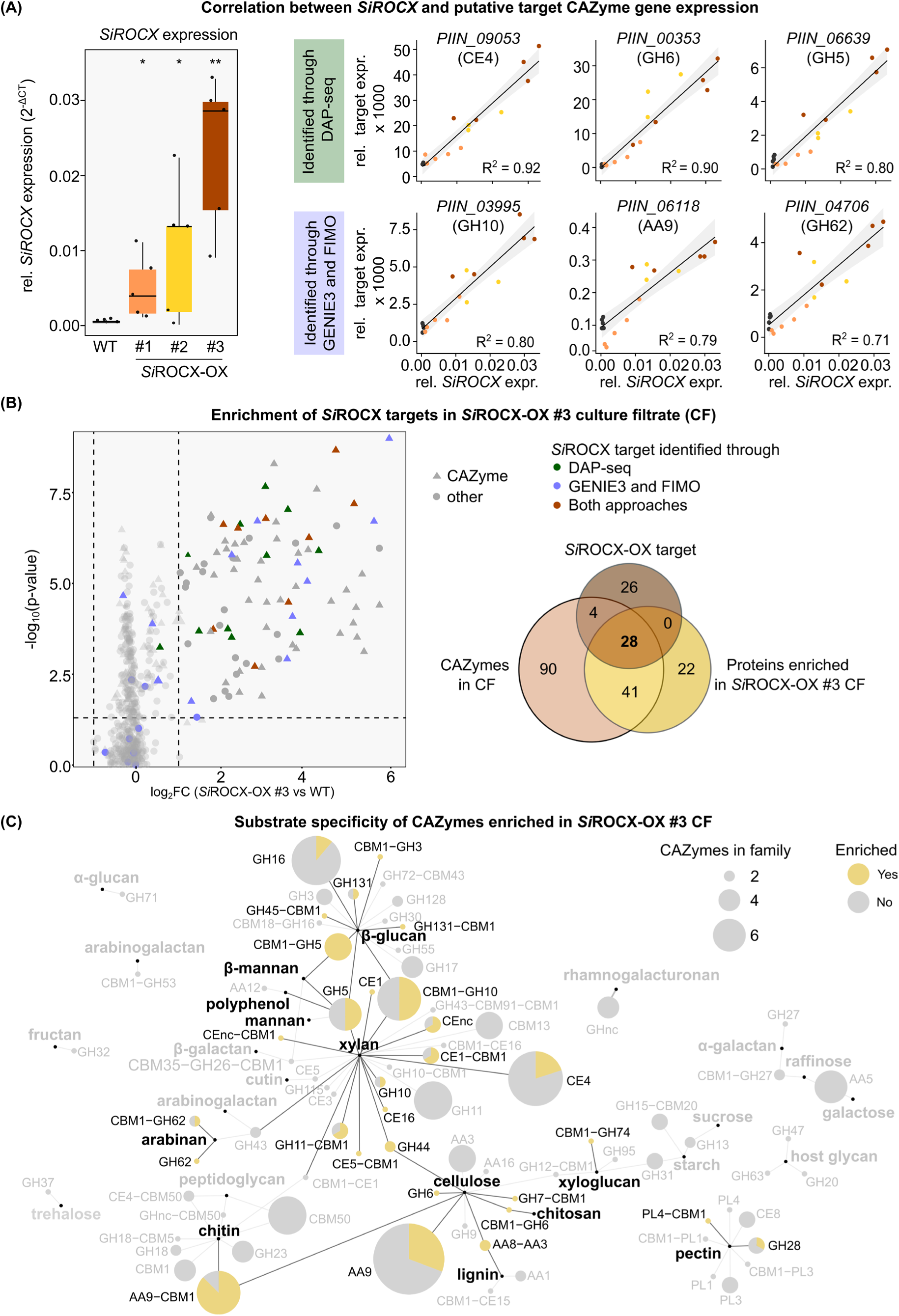
*SiROCX* overexpression increases secretion of CAZymes targeting cellulose and xylan in axenic culture. **A)** Wild-type *Serendipita indica* (WT) and three independent transformants (*Si*ROCX-OX #1–#3) expressing *SiROCX* under control of the inducible *Si*FGB1 promoter were grown in liquid CM medium. *SiROCX* expression correlated with expression of representative *Si*ROCX target genes as determined by qRT-PCR (n = 5 per transformant). Scatterplots show correlations between *SiROCX* expression and expression of selected CAZyme targets. Significant differences relative to WT are indicated by asterisks (* p < 0.05; ** p < 0.01, Student’s t-test). **B)** Proteins precipitated from culture filtrates were analyzed by mass spectrometry to identify proteins enriched in the *Si*ROCX-OX #3 secretome. The volcano plot shows differential protein abundance between WT and *Si*ROCX-OX #3. CAZymes are shown as triangles and other proteins as circles. Dashed lines indicate thresholds of |log₂FC| ≥ 1 and –log10(p-value) ≥ 1.3. Proteins significantly enriched in the *Si*ROCX-OX #3 strain relative to WT are highlighted. **C)** Network of secreted *S. indica* CAZymes annotated by predicted substrate class and domain composition. Node size reflects the number of CAZymes within each family. Pie charts indicate the fraction of CAZymes significantly enriched in the *Si*ROCX-OX#3 culture filtrate relative to WT (log₂FC ≥ 1, p < 0.05; yellow segments). CAZyme annotations are based on CAZyDB (http://www.cazy.org/), dbCAN3-HMMER, dbCAN3-sub 4-5, and manual subfamily curation (Brands et al., 2025).

We next asked whether *SiROCX* overexpression alters the secreted enzyme repertoire. Proteins were precipitated from culture filtrates (CFs) of WT and *Si*ROCX-OX #3 and analyzed by mass spectrometry. In total, 521 proteins were detected across both CFs (Fig. 2B, Supplementary Table S5). No proteins were significantly enriched in WT, whereas 91 proteins were significantly enriched in *SiROCX*-OX #3 (>2-fold change, p < 0.05). Among these were the acetyl-esterase *Si*AXE (PIIN_07653) and the endo-xylanase *Si*GH11 (PIIN_05889), two CAZymes previously shown to be co-induced during monocot colonization and to act sequentially in xylan deconstruction. *Si*GH11 releases *O*-acetylated xylo-oligosaccharides that are subsequently deacetylated by *Si*AXE, facilitating downstream hydrolysis and limiting generation of immunogenic fragments (Brands et al., 2025). Motif scanning identified a *Si*ROCX binding site in the *Si*GH11 promoter, whereas *Si*AXE was not predicted as a direct target, consistent with indirect regulation.

Across the secretome proteomics dataset, we detected 163 CAZymes, of which 69 (42%) were significantly enriched in the *Si*ROCX overexpressor relative to WT, with the strongest representation from AA9 (n=11), GH10 (n=9), and GH5 (n=9) families. These enriched CAZymes included five of the six qRT-PCR–validated targets (PIIN_00353, PIIN_06118, PIIN_04706, PIIN_09053, PIIN_03995). Overall, 28 of 69 enriched CAZymes (40%) were classified as direct *Si*ROCX targets based on DAP-seq peaks and/or motif-based prediction. The remaining enriched CAZymes likely reflect indirect regulation, potentially *via* secondary regulators, consistent with *Si*ROCX binding sites upstream of several transcription factors, including the M4 TF PIIN_09558 and the M7 TF PIIN_07406.

### *Si*ROCX-driven secretome deconstructs native barley root cell walls

As *Si*ROCX targets CAZymes with diverse predicted substrate specificities (Fig. 2C) that accumulate in the culture filtrates of the overexpression strains, we next characterized the enzymatic activities present in these filtrates. We assayed hydrolytic activity against a panel of artificial substrates, including azo-xylan (Fig. 3A), para-nitrophenyl (pNP) acetate (pNP-C2), and the long-chain ester pNP-palmitate (pNP-C16) (Fig. 3B). Alongside WT culture filtrates, we included filtrates from strains expressing GFP or the acetyl-xylan esterase *Si*AXE under the *Si*FGB1 promoter as controls (Brands et al., 2025). Purified *Si*GH11 and the promiscuous lipase *Si*PLL (PIIN_06162) served as positive controls for endo-xylanase and lipase activity, respectively.

**Fig. 3.**
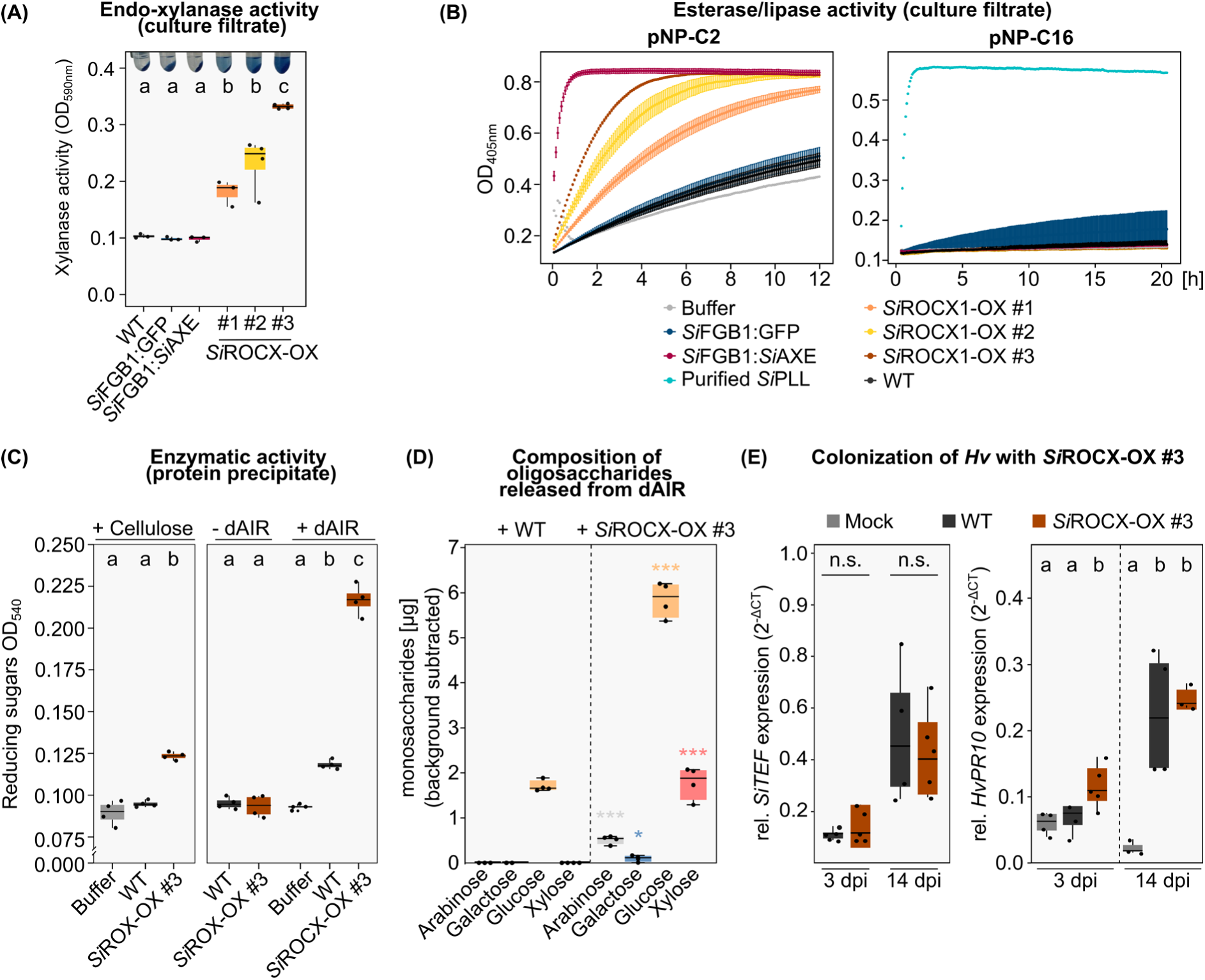
*SiROCX* overexpression enhances xylanase and cellulase-associated activities in axenic culture and increases barley wall sugar release. Transformants expressing *SiROCX* under the inducible *Si*FGB1 promoter (*Si*ROCX-OX) were grown in liquid CM medium for seven days, crushed, allowed to regenerate for three days, and cell-free culture filtrates (CFs) were harvested. CFs from strains expressing GFP or the acetylxylan esterase *Si*AXE (PIIN_07653) under the control of the *Si*FGB1 promoter served as controls. **A)** Endo-xylanase activity in CFs measured on azo-xylan (OD_590_) for WT, *Si*FGB1:GFP, *Si*FGB1:*Si*AXE, and three independent *Si*ROCX-OX strains (#1–#3). **B)** Esterase/lipase activity kinetics in CFs on pNP substrates: pNP-C2 (esterase; left) and pNP-C16 (lipase; right), monitored as OD_405_ over time. Buffer control and purified *Si*PLL (PIIN_06162) are included as negative and positive controls, respectively. **C)** Reducing sugar release (DNS assay; OD_540_) from cellulose (left) and from delignified alcohol-insoluble residue (dAIR) prepared from barley roots (right) using proteins precipitated from CFs. “−dAIR” indicates a no-substrate control. **D)** Monosaccharides (arabinose, galactose, glucose, xylose) were measured after chemical hydrolysis of oligosaccharides in the supernatant of barley dAIR after digestion with precipitated secreted proteins from WT or *Si*ROCX-OX #3 CFs. Background values from dAIR incubated with buffer were subtracted. Asterisks indicate two-tailed students t-test with Holm–Šídák correction (*p adj. ≤ 0.05, *** p adj. ≤ 0.001) of the respective monosaccharide (indicated by the same color) between WT and *Si*ROCX-OX #3. **E)** Barley (*Hordeum vulgare*) colonization with *Si*ROCX-OX #3. Fungal colonization level quantified by qRT-PCR (left) and host defence marker *HvPR10* expression (right) at 3 and 14 dpi compared to mock and WT. Boxplots show individual biological replicates (points). Different letters indicate significant differences (one-way ANOVA with Tukey’s post hoc test, p ≤ 0.05); n.s., not significant.

All three independent *Si*ROCX-OX culture filtrates exhibited a significant increase in endo-xylanase activity compared to WT and control strains, with the magnitude of the activity increase correlating with *SiROCX* expression levels (Fig. 3A). A similar pattern was observed for acetyl-esterase activity with pNP-C2 but not lipase activity with pNP-C16 (Fig. 3B). Together, these data indicate that *SiROCX* overexpression increases secreted enzymatic activities consistent with plant cell wall polysaccharide deconstruction.

To test whether these activities translate into enhanced degradation of native monocot cell walls, we precipitated proteins from culture filtrates of the strongest *Si*ROCX overexpressor (*Si*ROCX-OX #3) and incubated them with microcrystalline cellulose or delignified alcohol-insoluble residue (dAIR) prepared from barley roots. Enzymatic activity was quantified by measuring released reducing sugars in the supernatant (Fig. 3C). On cellulose, only proteins from the *Si*ROCX-OX culture filtrate released detectable sugars, whereas protein preparations from both WT and *Si*ROCX-OX filtrates released sugars from the natural dAIR substrate. However, the *Si*ROCX-OX sample liberated nearly fivefold more sugars from dAIR, indicating a marked increase in cell wall–degrading capacity.

The monosaccharide composition of oligosaccharides released from dAIR degradation was analyzed after acid hydrolysis. The most abundant monosaccharides in *Si*ROCX-OX #3–incubated samples were glucose and xylose, while smaller amounts of arabinose and galactose were detected (Fig. 3D). This indicates that enzymes secreted by the *Si*ROCX-OX #3 strain released a broader spectrum of wall-derived oligosaccharides than WT, likely originating from xylooligosaccharides (XOS), cellooligosaccharides (COS), and mixed-linkage glucan (MLG) fragments, consistent with coordinated hemicellulose and cellulose deconstruction. Collectively, these results establish *Si*ROCX as a driver of a multi-enzyme secretome that synergistically couples xylan deconstruction (*via* debranching, deacetylation, and hydrolysis) with cellulose degradation on native barley root cell walls.

This enhanced wall-deconstruction capacity aligns with observations *in planta*. Unlike WT *Si*, *Si*ROCX-OX #3 triggered expression of the immune marker gene *HvPR10* during early barley colonization (3 dpi), despite comparable fungal biomass (Fig. 3E), consistent with premature or excessive CAZyme activity. These findings indicate that precise temporal activation of *Si*ROCX is required to enable nutrient acquisition from accessible cell wall substrates while avoiding immune activation during biotrophic accommodation.

### *Si*ROCX exhibits host-, stage-, and cell wall–derived cue–specific regulation

Given the monocot-specific induction of *Si*ROCX-regulated CAZymes and their cell wall–degrading activities (Fig. 1; Brands et al., 2025), we next examined whether *Si*ROCX itself is subject to host- and stage-dependent regulation during root colonization. Colonization assays with WT *S. indica* revealed strong induction of *Si*ROCX transcripts during late-stage barley colonization (14 dpi), whereas expression remained low during colonization of *Arabidopsis thaliana*. This host-dependent expression pattern is consistent with the association of *Si*ROCX with the monocot-induced WGCNA Module M7 (Fig. 1A; Fig. 4A).

**Fig. 4.**
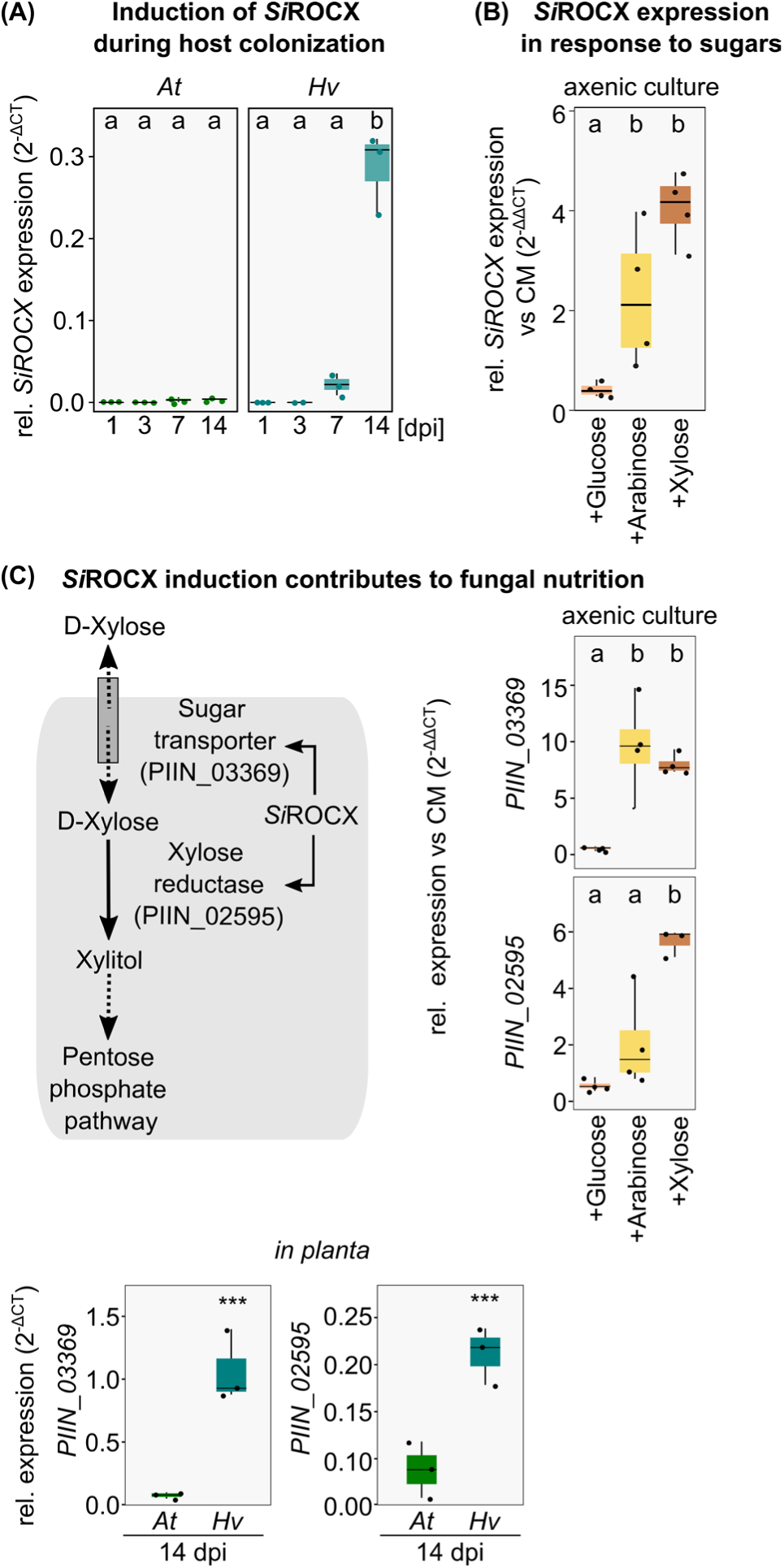
*SiROCX* is induced during late-stage monocot but not dicot colonization and in response to different cell wall stimuli. **A)** Barley (*Hordeum vulgare*, *Hv*; monocot) and *Arabidopsis thaliana* (*At*; dicot) roots were inoculated with *Si* and sampled at different colonization stages. *SiROCX* expression was measured by qRT-PCR relative to the fungal housekeeping gene *SiTEF* and shows strong induction during late-stage colonization of *Hv* but not *At*. **B)** qRT-PCR analysis of *SiROCX* expression in axenic culture medium supplemented with glucose, arabinose, or xylose shows induction by xylose and arabinose. *SiROCX* expression was normalized to expression in medium without monosaccharides (CM medium) **C)** Model linking xylose utilization to *Si*ROCX-dependent regulation of a sugar transporter (PIIN_03369) and xylose reductase (PIIN_02595) in the pentose catabolic pathway. Expression of PIIN_03369 and PIIN_02595 was quantified by qRT-PCR relative to fungal *SiTEF* during late-stage colonization (14 dpi) of *Hv* and *At* and in CM medium supplemented with glucose, arabinose, or xylose. Expression in medium with monosaccharides was normalized to expression in medium without monosaccharides (CM medium). Boxplots show biological replicates (points). Different letters indicate significant differences (one-way ANOVA with Tukey’s post hoc test, p ≤ 0.05). Asterisks indicate significant differences with two-tailed student t-test in pairwise comparisons as shown (* p < 0.05; ** p < 0.01; *** p < 0.001).

**Fig. 5.**
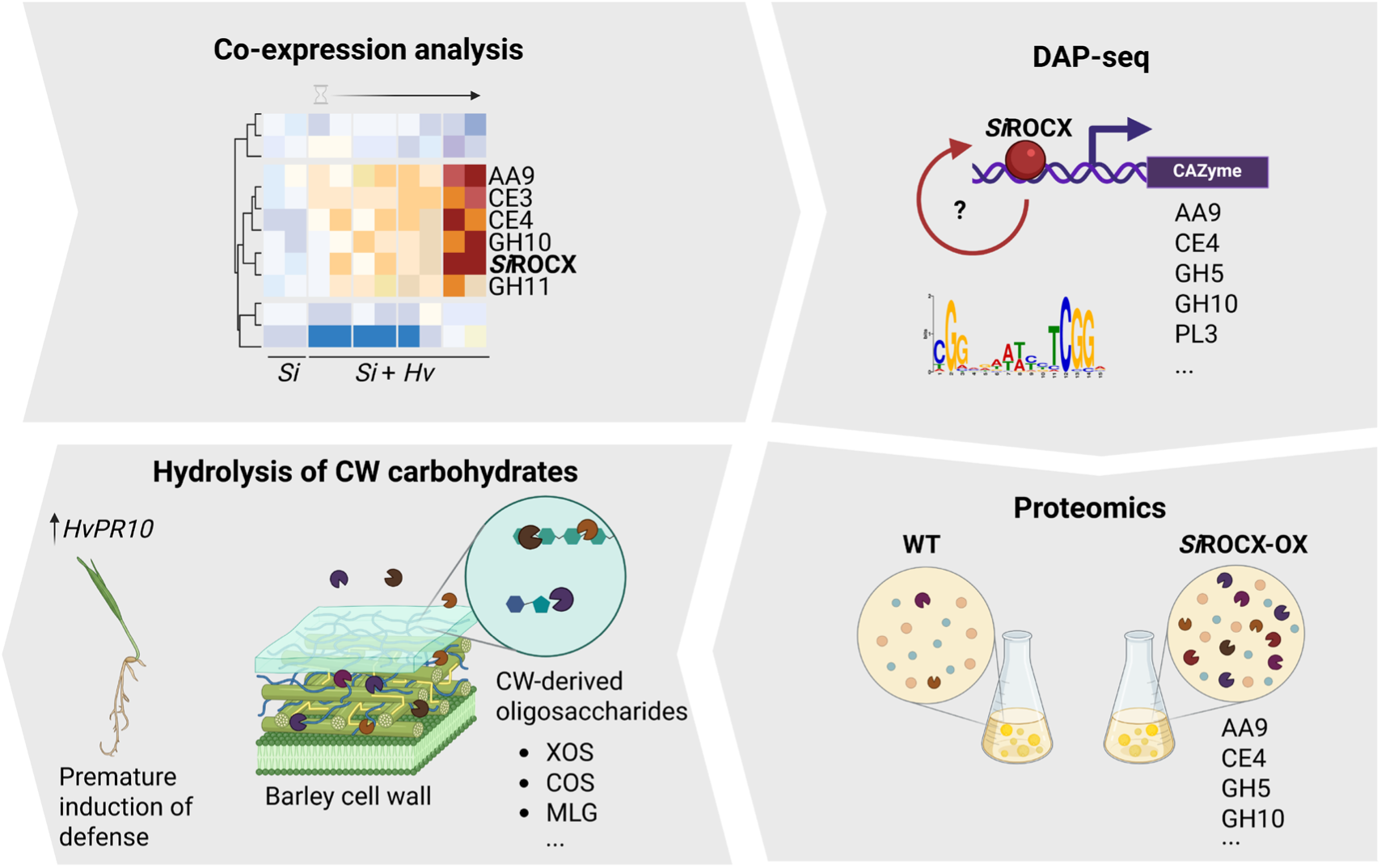
Model of *Si*ROCX-driven monocot cell wall deconstruction during late-stage colonization by *Serendipita indica*. Co-expression analysis during monocot root colonization identifies a CAZyme-enriched module associated with *Si*ROCX. DAP-seq defines the *Si*ROCX binding motif and a CAZyme-biased regulon. This motif is also present upstream of *Si*ROCX itself suggesting autoregulation. Inducible *SiROCX* overexpression increases secretion of xylan- and cellulose-active enzymes, and secreted proteins precipitated from culture filtrate release oligosaccharides that mainly consist of xylose and glucose, likely originating from xylooligosaccharides (XOS), cellooligosaccharides (COS), and mixed-linkage glucan fragments (MLG) from barley root cell wall material. Together, these data support *Si*ROCX as a central regulator coordinating a monocot-adapted cell wall–degrading program whose timing is important for symbiotic compatibility. Figure created with BioRender.

To identify potential plant-derived cues, we quantified *SiROCX* expression in WT fungi grown in axenic culture in liquid CM medium supplemented with glucose, arabinose, or xylose. These sugars served as proxies for the final products generated by complete degradation of the monocot host cell wall polysaccharides cellulose, MLG, and xylan. While expression remained low in glucose-supplemented medium, arabinose and xylose caused a two- to threefold increase in *SiROCX* expression (Fig. 4B).

Next, we examined induction of PIIN_03369 and PIIN_02595, two M7 genes and *Si*ROCX targets putatively involved in the uptake and catabolism of pentose sugars derived from the host cell wall. PIIN_03369, encoding a sugar transporter, was induced eight- to tenfold by arabinose and xylose, whereas PIIN_02595, encoding a xylose reductase, was induced approximately sixfold by xylose; expression remained low in glucose for both genes (Fig. 4C). Both genes also showed late-stage induction during barley colonization (14 dpi), but not in *A. thaliana* (Fig. 4C), consistent with the expression pattern observed for *SiROCX*. Because arabinose and xylose are principal constituents of monocot hemicellulose, these findings support a model in which their availability contributes to host-specific activation of *Si*ROCX during barley colonization.

## Discussion

Successful colonization of plant roots by beneficial fungi requires a balance between host cell wall remodeling for nutrient access and immune compatibility. The plant cell wall serves both as a structural barrier rich in polysaccharides and as an immunologically active interface; microbial CAZymes must therefore be precisely coordinated to breach this barrier while minimizing the release of damage-associated molecular patterns (DAMPs) and activation of host defences. Our study identifies the basidiomycete-specific transcription factor *Si*ROCX as a key regulator of a monocot-adapted CAZyme program in *S. indica*, revealing a regulatory layer that links host-derived signals to controlled cell wall remodeling during symbiotic colonization.

Recent work demonstrated that *S. indica* deploys a cooperative enzymatic module for the deconstruction of acetylated xylan, the dominant hemicellulose in grass cell walls. This module includes the endo-xylanase *Si*GH11, which releases acetylated XOS, and the acetyl-xylan esterase *Si*AXE, which subsequently removes acetyl groups from soluble oligomers to facilitate further hydrolysis and reduce the accumulation of immunogenic intermediates. The sequential activity of these enzymes allows efficient degradation of monocot hemicellulose while simultaneously limiting immune activation, illustrating how enzymatic specialization contributes to immune-compatible symbiosis. However, while the biochemical functions of these enzymes have been elucidated, the upstream regulatory mechanisms controlling their coordinated expression during root colonization remained unknown.

Our findings provide evidence that *Si*ROCX functions as a transcriptional activator of this monocot-specific cell wall–degrading program. Integrating transcriptomics, DAP-seq, and secretome proteomics revealed that *Si*ROCX directly or indirectly regulates a broad set of CAZymes targeting xylan and cellulose, including enzymes from the GH, CE, and AA families that together constitute a complex extracellular degradation network. Overexpression of *SiROCX* markedly increased the secretion of these enzymes and enhanced hydrolytic activity on both artificial substrates and native barley root cell walls, demonstrating that *SiROCX* expression is sufficient to drive a functional cell wall–degrading secretome. These findings suggest that the enzymatic modules described previously, such as the *Si*GH11–*Si*AXE–GH43 cascade, are embedded within a larger transcriptional program coordinated by *Si*ROCX.

Importantly, this CAZyme program is tightly restricted to later stages of colonization. During the early biotrophic phase of the interaction, extensive host cell wall degradation would likely trigger plant immune responses and compromise intracellular accommodation. Consistent with this, both CAZyme expression and *Si*ROCX activity remain low during early colonization, when fungal biomass is still limited and host cells remain alive. Previous studies have shown that *S. indica* undergoes a biphasic colonization strategy, transitioning from an initial biotrophic phase to a later saprotrophic-like phase associated with localized host cell death (Lahrmann et al., 2015; Eichfeld et al., 2024). Our results suggest that activation of the *Si*ROCX regulon contributes to this transition by promoting controlled cell wall remodeling once symbiotic compatibility has been established. One mechanism that may contribute to this temporal control is carbon catabolite repression (CCR). During early colonization, the fungus likely relies on readily available host-derived sugars such as glucose, which repress the expression of plant cell wall–degrading enzymes across fungi (Adnan et al., 2017). As simple sugars become depleted or hemicellulose-derived sugars such as arabinose and xylose accumulate, CCR may be relieved, allowing activation of the *Si*ROCX regulon. In this model, host-derived sugars link fungal metabolic state to CAZyme deployment, enabling the transition from biotrophic accommodation to late-stage cell wall remodeling while maintaining overall tissue-level compatibility.

The discovery of *Si*ROCX also places the regulation of plant biomass degradation in a broader evolutionary context. In ascomycete fungi, CAZyme expression is controlled by well-characterized regulators such as XlnR, CLR-1, and CLR-2. In contrast, basidiomycete regulatory networks have remained poorly understood despite their extensive lignocellulose-degrading capabilities. The transcription factor Roc1, identified in the wood-decaying fungus *Schizophyllum commune*, represents the first basidiomycete-specific regulator of cellulase expression (Marian et al., 2022). Roc1 directly controls genes encoding lignocellulose-degrading enzymes and is essential for growth on cellulose-containing substrates.

Phylogenetic analyses indicate that *Si*ROCX belongs to this Roc1/ROCX lineage, suggesting that these regulators represent an ancient basidiomycete-specific transcriptional module controlling plant biomass degradation. In saprotrophic fungi such as *S. commune*, this regulatory system orchestrates the decomposition of lignocellulosic substrates in dead plant material. In contrast, in the mutualistic endophyte *S. indica*, the same regulatory framework appears to have been repurposed to enable controlled remodeling of living host cell walls.

This functional repurposing likely reflects the evolutionary transition from saprotrophic ancestors to symbiotic lifestyles within the Sebacinales. *S. indica* retains a large repertoire of CAZymes reminiscent of decomposer fungi, yet their expression is tightly regulated and restricted to specific colonization stages and host contexts. Our data indicate that *Si*ROCX activation occurs primarily during late-stage colonization of monocot roots and is responsive to pentose sugars such as arabinose and xylose, which are derived from hemicellulose breakdown. Because monocot cell walls are particularly enriched in glucuronoarabinoxylan, this regulatory module may be especially advantageous during colonization of grass hosts. This suggests that cell wall–derived sugars act as environmental cues linking substrate availability to transcriptional activation of the CAZyme program.

Together, these findings reveal how transcriptional regulation and specialized enzymatic modules cooperate to enable immune-compatible host cell wall remodeling during fungal symbiosis. At the regulatory level, the basidiomycete-specific transcription factor *Si*ROCX activates a coordinated CAZyme secretome. At the enzymatic level, specialized modules such as the *Si*GH11–*Si*AXE cascade process host polysaccharides into non-immunogenic sugars. This multi-layered strategy allows *S. indica* to access host-derived carbon while maintaining a compatible interaction with the plant. More broadly, these findings illustrate how ancestral lignocellulose-degrading pathways have been repurposed during fungal evolution to support symbiotic lifestyles, with Roc1/ROCX transcription factors emerging as key regulators controlling the deployment of this program across ecological contexts ranging from saprotrophy to symbiosis.

## Material and Methods

### Plant and fungal materials

The root endophyte *Serendipita indica* (*Si*; DSM 11827, Deutsche Sammlung von Mikroorganismen und Zellkulturen, Braunschweig, Germany) was used as a model fungal symbiont. *Hordeum vulgare* L. cv. Golden Promise (*Hv*), *Brachypodium distachyon* (*Bd*, Bd21-3) and *Arabidopsis thaliana* Col-0 (*At*) were employed as host plants for colonization studies.

### Fungal growth and chlamydospore isolation

*Si* was cultured on CM medium (Hilbert et al., 2012) composed of 20 g L^-1^ glucose, 2 g L^-1^ peptone, 1 g L^-1^ yeast extract, 1 g L^-1^ caseine hydrolysate and 1 ml L^-1^ microelement solution (6 g MnCl_2_·4H_2_O, 1.5 g H_3_BO_3_, 2.65 g ZnSO_4_ ·7H_2_O, 750 mg Kl, 2.4 mg Na_2_MO_4_·2H_2_O, 130 mg CuSO_4_·5H_2_O and 50 ml L^-1^ salt solution (120 g NaNO_3_, 10.4 g KCl, 10.4 g MgSO_4_·7H_2_O, 30.4 g KH_2_PO_4_). The medium was solidified with agar (15 g L^-1^). Cultures were grown at 28 °C in the dark for four weeks.

For chlamydospore isolation, 5 mL Tween water (0.002 % v/v Tween-20 in MQ-H_2_O) were added to the surface of a four-week-old CM agar plate containing actively growing mycelium. Spores were gently dislodged with a sterile scalpel, and the resulting suspension was filtered through Miracloth (22-25 µm pore size) to remove mycelium fragments. The filtrate was collected in a 50 mL centrifuge tubes and washed three times with Tween water by centrifugation at 3,350 g for 7 min. After the final wash, the spores were resuspended in MQ-H_2_O. The resulting spore suspension was either directly used to inoculate 100 mL of CM liquid culture (for axenic fungal cultures) or diluted to 500.000 mL^-1^ for plant inoculation experiments.

### RNA-seq and differential gene expression analysis

Time-resolved RNA-seq data of *Serendipita indica* (*Si*) colonizing the roots of *Hordeum vulgare* (*Hv*) or *Brachypodium distachyon* (*Bd*) was obtained as previously described (Eichfeld et al., 2024). The raw data of the RNA-seq experiments performed at the US Department of Energy Joint Genome Institute (JGI) under a project proposal (ID: 505829; Zuccaro, 2020) can be downloaded from the NCBI BioProject database (https://www.ncbi.nlm.nih.gov/bioproject/) under the accession numbers PRJNA1052152-PRJNA1052163 (*Si*), PRJNA1052227-PRJNA1052238 (*SiBd*) and PRJNA1052254-PRJNA1052262 (*SiHv*). Significant differentially expressed genes (*Si* vs. *SiBd* or *SiHv*) were identified for all timepoints (1, 3, 6 and 10 days post inoculation) using the DESeq2 package version 1.38.3 in R (Love et al., 2014). For the generation of heatmaps, log_2_FC values were visualized using the pheatmap package version 1.0.12 (Kolde, 2025).

## gDNA extraction2 from fungal mycelium

For genomic DNA extraction, *Si* mycelium was grown in liquid CM medium for seven days, crushed and after three days of recovery harvested through a Miracloth (22-25 µm pore size) filter. The mycelium was flash frozen in liquid nitrogen and ground to a fine powder. 500 mg of powder were homogenized in 4 mL extraction buffer (250 mM NaCl, 25 mM EDTA, 0.5 % (w/v) SDS, 250 mM Tris-HCl, pH 7.5). Proteins and debris were removed by potassium acetate precipitation. In brief, 1.5 mL 3 M KAc (pH 5.0) were added to each sample and after centrifugation at 17,700 g for 5 min, the debris-free supernatant was mixed with 5,5 mL 100 % (v/v) isopropanol and incubated ON at 4 °C. The next day, the samples were centrifuged at 17,700 g for 10 min and the resulting pellet was washed twice with 5 mL 70 % (v/v) ethanol through centrifugation at 17,700 g for 3 min. Subsequently, the pellet was dried at room temperature and resuspended in 500 µL water. The pellet was further purified through sequential phenol–chloroform and chloroform extractions, followed by isopropanol precipitation. For this purpose, the sample was mixed with 1 V phenol:chloroform:isoamylalcohol (25:24:1), incubated on ice for 30 min, and centrifuged at 13,500 g for 10 min. The aqueous phase was transferred to a fresh tube, extracted with 1 V chloroform:isoamyl alcohol (24:1), and centrifuged under the same conditions. DNA was precipitated from the recovered aqueous phase by adding 1 V of 100 % isopropanol and 1/10 V of 3 M sodium acetate (pH 6.0), followed by incubation on ice for 30 min and centrifugation at 17,700 × g for 10 min. The resulting DNA pellet was washed twice with 70% ethanol, air-dried at room temperature, and resuspended in 100 µL of water. RNA contamination was removed by treatment with 10 µg RNase A for 1 h at 37 °C, and DNA was re-precipitated by chloroform isoamylalcohol extraction as described above.

### DNA affinity purification sequencing (DAP-seq)

Genes encoding *Si* transcription factors (as annotated in Mycocosm, Grigoriev et al. (2014)) were codon-refactored using the BOOST software suite with an *Escherichia coli* (*E. coli*) codon usage table and the balanced codon algorithm (Oberortner et al., 2017). Synthetic gene fragments were obtained from Twist Biosciences and designed with 30 bp vector-compatible overlaps to enable assembly as N-terminal HALO-tag fusions. Fragments were PCR-amplified using KAPA DNA polymerase (Roche), purified in a 384-well plate format using the Wizard SV PCR Purification kit (Promega), and assembled into the pIX-HALO-ccdB vector by Gibson assembly (NEBuilder HiFi DNA Assembly Master Mix, NEB) (Gibson et al., 2009). Assembly reactions were transformed into *E. coli* TOP10 (Thermo Fisher), and individual colonies were screened by PCR and verified by sequencing on the Pacific Biosciences Sequel II and IIe platforms using custom analysis pipelines.

MultiDAP-seq experiments were conducted using the protocol described previously (Baumgart et al., 2021) with minor modifications for eukaryotic transcription factors (Baumgart et al., 2025). Genomic DNA from *Si* was sheared to an average size of 150 bp with a Covaris LE220-Plus focused-ultrasonicator (Covaris), followed by library preparation using the KAPA HyperPrep kit (Roche) with the same adapters as described previously (Baumgart et al., 2021) to ligate a unique i7 index primer to each genomic DNA library. Upstream of the DAP-seq assay, genomic DNA libraries were PCR amplified for 10 cycles to remove DNA modifications. Halo-tagged transcription factor protein expression templates were PCR amplified directly from *E. coli* glycerol stocks using the primers pIX-Halo-T7-fwd (5′-GTGAATTGTAATACGACTCACTATAGGG) and pIX-Halo-AfterPolyA-rev (5′-CAAGGGGTTATGCTAGTTATTGCTC), purified using SPRI beads, and the correct size of each was verified using a Fragment Analyzer (Agilent Technologies). At least 500 ng of each linear PCR product was used for *in vitro* protein expression using the TnT T7 Quick for PCR DNA kit (Promega). Each DAP-seq reaction was run with 100 uL expressed protein, a mixture of the two previously prepared fragment libraries, and 5000 ng salmon sperm DNA to reduce non-specific binding. Input genomic DNA library amounts were titrated on the basis of genome size using 20 ng each reaction. The binding incubation, washes, and final library amplification and purification were performed as in the multiDAP-seq protocol described previously (Baumgart et al., 2025). The final DAP-seq libraries were sequenced with a NovaSeq X Plus (Illumina), targeting 2-3 million 2×150 reads per sample.

Sequenced reads were adapter trimmed and quality filtered using BBTools version 38.90 (https://sourceforge.net/projects/bbmap/) using the following options: k=21 mink=11 ktrim=r tbo tpe qtrim=r trimq=6 maq=10. Filtered reads were aligned to the reference genome using bowtie2 (Langmead and Salzberg, 2012) version 2.4.2 with the options --no-mixed --no-discordant. Peaks were called using MACS3 version 3.0.0a6 (Zhang et al., 2008) using combined negative control samples with mock protein expression as the background file, and the following options: --call-summits --keep-dup 1, and --gsize with the total reference genome size. Sequences representing up to the 100 strongest peaks, as scored by signal value in column 7 of the narrowPeak files, were extracted using a custom python script. These peak sequences were used to generate motifs using MEME (MEME Suite v.5.3.0) (Bailey et al., 2009) with a zero-order background file generated from the reference genome, and the following additional options: -dna -revcomp -mod anr -nmotifs 2 -minw 8 -maxw 32. Motif calling was run twice for each dataset, once with the entire peak regions and once with only the regions +/- 30 bp flanking the summit location of each peak. Peaks were assigned to putatively regulated genes with BEDTools version 2.30.0 (Quinlan and Hall, 2010), using bedtools subtract to filter for intergenic peaks, followed by bedtools closest to assign each peak to up to two genes immediately adjacent to and downstream of the peak.

In total, 62 of 291 successfully synthesized TFs (21 %) passed quality control (≥100,000 filtered aligned fragments, FRIP score ≥0.01, at least one peak with a signal intensity ≥15-fold over the background) and yielded high-confidence TF binding motifs.

### Chord diagrams

The module membership and substrate specificities as described in Brands et al. (2025) were visualized using the chordDiagram() function of the circlize package for circular representation in R (Gu et al., 2014).

### *In silico* prediction of transcription factor targets

RNA-seq data was analysed with a WGCNA approach (Langfelder and Horvath, 2008) in R as described previously in Brands et al. (2025). In brief, raw counts were normalized using the a variance stabilizing vst() transformation from the DESeq2 package version 1.38.3 in R (Love et al., 2014). Subsequently, samples with < 500.000 reads and genes with < 10 raw read counts in >10 % were excluded from further analysis. In consequence, expression values for 10,658 of 11,767 annotated *Si* genes (90 %) were used as input for the WGCNA (version 1.72). The blockwiseModules() function was run with the following parameters: power = 12, networkType = “signed hybrid”, minModuleSize = 100, mergeCutHeight = 0.15. In total, 8,657 genes were assigned to seven different modules whereas 2,011 genes remained unassigned (M0). Compared to the whole *Si* genome as background, two modules (M4 and M7) were >2-fold enriched for genes encoding carbohydrate active enzymes (CAZymes) as annotated in Mycocosm (https://mycocosm.jgi.doe.gov/mycocosm/home). To predict target genes of transcription factors (TFs) with high membership scores (>0.75) to M4 and M7, a complementary network analysis using the GENIE3 package in R was conducted with default settings (Huynh-Thu et al., 2010). The analysis included 456 *Si* TFs annotated in Mycocosm, along with the DNA-binding protein PIIN_08210, which harbours a DNA-binding domain characteristic of a fungal TF family (IPR003163). For each TF, predicted target genes with an edge weight (connectivity) > 0.01 were extracted for downstream analysis.

The TF binding motifs identified through DAP-seq were used as input for FIMO to scan the promoter regions of *Si* genes (defined as 1,000 bp regions upstream of the start codon excluding exons) for potential TF binding sites (Grant et al., 2011).

The information obtained from GENIE3 and the *in silico* search for TF binding motifs was combined to expand the list of potential TF targets. Genes were classified as TF targets if they harboured a binding motif and were linked to the respective TF in the GENIE3 network.

### Overexpression of *SiROCX*

The full-length coding sequence of *SiROCX* as annotated in the genome assembly GCA_000313545 (*SiROCX*_1-480_) was amplified by PCR from *Si* cDNA (primers listed in Supplementary Table S6). The PCR product was cloned into the K100 expression vector (Wawra et al., 2016) under the control of either the *Si*FGB1 or *Sv*TEF promotor using Gibson assembly (NEB; #E2611). This construct added a C-terminal HA- and a His_6_-tag to *SiROCX*.

Transformation of *Si* was carried out as previously described (Wawra et al., 2016). In brief, *Si* mycelium grown in liquid culture for seven days was harvested through Miracloth (22-25 µm pore size) and washed with MQ-H_2_O containing 0.9 % NaCl. The mycelium was then resuspended in 100 mL fresh CM medium and crushed with a blender. The resulting fragments were incubated for 2 days at 28 °C while shaking at 130 rpm to allow regeneration.

The regenerated mycelium was filtered again through Miracloth, washed with 0.9 % NaCl solution, and subjected to enzymatic digestion in 20 mL sterile SMC medium (1.33 M sorbitol, 50 mM CaCl_2_, 20 mM MES, pH 5.8) supplemented with 20 g L^-1^ *Trichoderma harzianum* lysing enzymes (Sigma, #L1412). After 1-2 h at 32 °C, the protoplasts were filtered through Miracloth and collected in a 50 mL centrifugation tube. The enzymatic reaction was stopped by adding 10 mL ice-cold STC buffer (50 mM CaCl_2_, 1M sorbitol, 10 mM Tris-HCl, pH 7.5).

Protoplasts were pelleted by centrifugation 4,000 rpm and 4 °C for 10 minutes and then resuspended in 1 mL ice-cold STC solution. This washing step was repeated twice. For transformation, 70 µL protoplast suspension were mixed with 10 µg linearized plasmid DNA and 1 µl heparin (15 mg mL^-1^). After 10 min on ice, 0.5 mL sterile ice-cold STC solution supplemented with 40 % PEG 3350 was added. The mixture was incubated for 15 min on ice, then transferred to a 15 mL reaction tube containing 5 mL of hand-warm liquid top medium (7 g L^-1^ malt extract, 1 g L^-1^ pepton, 0.5 g L^-1^ yeast extract, 0.3 M sucrose, 0.6 % w/v agar). The mixture was plated onto bottom medium (7 g L^-1^ malt extract, 1 g L^-1^ pepton, 0.5 g L^-1^ yeast extract, 0.3 M sucrose, 1.2 % w/v agar) supplemented with 80 mg L^-1^ hygromycin for selection. Plates were incubated at 28 °C in the dark. Colonies appearing after 14 days were transferred to CM agar plates containing 80 µg mL^-1^ hygromycin for further propagation.

### Genotyping

To distinguish between dikaryotic and homokaryotic transformants, PCR amplification of multiallelic homeodomain (HD) genes PIIN_09977 (HD1.2) and PIIN_ 09916 (HD2.1) was performed (Zuccaro et al., 2011). Successful dikaryotic transformants were identified based on the presence of the expected transgene amplicon in combination with the correct HD gene pattern. Primer sequences are listed in Supplementary Table 1.

### Preparation of fungal culture filtrate and protein precipitate of secreted protein

To obtain metabolically active, rapidly growing *Si* mycelium, a 100 mL starter culture of CM medium was inoculated with chlamydospores and incubated for one week at 28°C while shaking at 120 rpm. The mycelium was harvested using Miracloth (22-25 µm pore size) and washed with sterile MQ-H2O containing 0.9 % NaCl. The washed mycelium was resuspended in 100 mL fresh CM medium and homogenized with a blender. The resulting mycelial fragments were incubated at 28 °C while shaking at 130 rpm to allow regeneration. After three days, the culture filtrate (CF) was separated from the mycelium using Miracloth (22-25 µm pore size). The mycelium was washed with 0.9 % NaCl, blotted dry on tissue paper and frozen in liquid nitrogen for protein or RNA extraction. The CF was kept at 4 °C for precipitation of secreted protein or enzymatic assays. PMSF was added to 1 a final concentration of 1 mM and the pH adjusted to 8 with 1M Tris buffer. Then, ammonium sulfate was added to a final amount of 40 % (w/v) and incubated at 1h with gentle agitation at 20°C. Protein were pelleted by centrifugation at 20900 *g*, the supernatant discarded and precipitated secreted protein resuspended in 20 mM Tris buffer pH=8.0 and used for enzymatic assays or protein mass spectrometry.

### Protein mass-spectrometry

Protein precipitates prepared from culture filtrate of WT and *Si*ROCX-OX #3 were analyzed by the CECAD Proteomics Facility on an Orbitrap Exploris 480 mass spectrometer (Thermo Scientific) coupled to a Vanquish neo in trap-and-elute setup (Thermo Scientific). Samples were loaded onto a 2 cm precolumn (Acclaim Pepmap 100, Thermo Scientific) with a flow of 10 µl/min of eluent A (0.1% formic acid) before in-line flushed onto an Auroa Ultimate column (25 cm length, 75 µm inner diameter, Ionopticks). Peptides were chromatographically separated with an initial flow rate of 400 nL/min and the following gradient: initial 4% B (0.1% formic acid in 80 % acetonitrile), up to 8.5 % in 4 min. Then, flow was reduced to 300 nl/min and B increased to 35% B in 75 min, and up to 98% solvent B within 1.0 min while again increasing the flow to 400 nl/min, followed by column wash and reequilibration to initial condition.

MS1 scans were acquired from 399 m/z to 1001 m/z at 30k resolution. Maximum injection time was set to 54 ms and the AGC target to 100%. MS2 scans ranged from 400 m/z to 1000 m/z and were acquired at 30 k resolution with a maximum injection time of 54 ms and an AGC target of 1000%. DIA scans covering the precursor range from 400 - 1000 m/z and were acquired in 30 x 20 m/z windows with an overlap of 1 m/z. All scans were stored as centroid.

Samples were analyzed in Spectronaut 19 (Biognosys) in directDIA mode using a *Serendipita indica* Ensmbl protein database. Search parameters were following recommended standard settings with minimum and maximum fragment filters set to six and 12, respectively. Afterwards, analysis of results was performed in Perseus 1.6.15 (Tyanova 2016). Low-intensity quantifications were converted to NaN, followed by log2 transformation of remaining LFQ values, filtering on data completeness in at least one replicate group and FDR-controlled T-tests between the two groups of interest.

### Esterase and lipase activity assays

Esterase and lipase activity was measured in protein extracts from axenically grown *Si* CF. *Para*-nitrophenyl (pNP-) acetate (Merck #46021) and pNP-palmitate (Merck, #N2752) were used as substrates. Stock solutions (1 mg mL^-1^) were prepared in ethanol by serial dilutions and pre-heated to 70 °C prior to pipetting to ensure complete solubilization. Recombinant *Si*PLL (Brands et al., 2025) served as positive control. Reaction mixtures were set up with 90 µL of assay buffer (10% [v/v] substrate, 1 % [v/v] Triton X-100, in 25 mM Tris-HCl, pH 7.5) and 10 µL of protein extract, CF or control protein solution. 100 µL of every reaction was transferred into a transparent 96-well microplate (Greiner). Plates were sealed and the absorbance at 405 nm (OD_405_) was measured at regular intervals overnight. Control reactions containing only buffer were included to account for non-enzymatic substrate hydrolysis.

### *In vitro* endo-xylanase activity assays

Endo-xylanase activity was measured in protein extracts from axenically grown *Si* CF. Recombinant *Si*GH11 (Brands et al., 2025) served as positive control. Assays were carried out with birchwood azo-xylan (Megazyme; #S-AXBP) as the substrate, following the manufacturer’s instructions. Briefly, 500 µL of azo-xylan solution (10 mg mL^-1^), 400 µL phosphate buffer (10 mM, pH 6.0) and 100 µL protein extract, CF or control enzyme were mixed and incubated for up to 24 h at RT. To quantify endo-xylanase activity, 200 µL aliquots were taken from the reaction mixture. To each aliquot, 500 µL of 95% (v/v) ethanol was added, followed by vortexing and incubation at 21 °C for 5 minutes. Samples were then centrifuged at 3,000 rpm for 10 minutes, and absorbance of the supernatant was measured at 590 nm (OD_590_) using a microplate reader.

### Generation of delignified alcohol insoluble residue (dAIR) from barley roots

Barley roots were harvested after 14 days of growth on 1/10 PNM medium, milled to a fine 648 powder using a Retsch MM400 mixer mill and lyophilized. Then, alcohol insoluble residue (AIR) was prepared as previously described (Kraemer et al., 2021). For delignification, AIR samples were incubated at 37 °C for 16 h in 0.5 % (g/v) ammonium oxalate, and 1 h at 85 °C in 11 % (v/v) peracetic acid (Gonçalves et al., 2008). The resulting delignified AIR (dAIR) was washed three times with water and twice with acetone. dAIR powder was resuspended in water to a concentration of 10 mg ml^-1^.

### Enzymatic hydrolysis of barley dAIR and cellulose

Ten mg ml^-1^ of barley dAIR slurry or microcrystalline cellulose (Merck; 9004-34-36) was incubated with protein precipitate of secreted protein (0.1 mg ml^-1^) from WT or *Si*ROCX-OX #3, for 16h. Then, soluble oligosaccharides were harvested by pelleting the insoluble material via centrifugation and the supernatant was harvested. The monosaccharide composition of barley dAIR-derived oligosaccharides was determined after acid hydrolysis using high-performance anion-exchange chromatography (HPAEC) as described previously (Brands et al., 2025). For quantification of sugars released from barley dAIR or cellulose, the dinitrosalicylic acid (DNSA) assay for reducing sugars was used. Supernatants were mixed with 1.5 volumes of DNSA reagent, incubated at 95°C for 10 minutes, cooled on ice, and absorbance was measured at 540 nm to determine the concentration of reducing sugars.

### Fungal inoculation and plant colonization assays

*Hv* seeds were surface sterilized by incubation in 6 % sodium-hypochlorite for 1 h, followed by six washes of 30 min each with sterile MQ-H_2_O. Seed coats were removed, and seeds were germinated on moist filter paper in the dark at 21 °C for three days. For fungal colonization, four germinated *Hv* seedlings were transferred to a sterile WECK jar containing 100 mL 1/10 PNM medium. Each jar was inoculated with 3 mL of *Si* chlamydospore suspension (500.000 spores ml^-1^). Plants were grown under long-day conditions (day/night cycle of 16/8 h, 22 °C/18 °C, light intensity of 108 μmol m-^2^s^-1^).

*At* seeds were surface sterilized by sequential incubation in 70 % ethanol for 10 min followed by 100 % ethanol for 7 min. After complete removal of ethanol, the seed sown on square plates filled with ½ MS medium (2.2 g L^-1^ MS salts [Duchefa, #M0245], 12 g L^-1^ Gelrite [Duchefa, #G1101], pH 5.7) supplemented with 10 g L^-1^ sucrose. Plates were stratified for three days at 4 °C in the dark, then transferred to short-day conditions (day/night cycle of 8/16 h, 21 °C, light intensity of 100-130 μmol m^-2^s^-1^) for germination. After fourteen days, the seedlings were transferred to fresh ½ MS plates without sucrose and inoculated with 1 mL of *Si* spore suspension.

For both hosts, roots were harvested 1-, 3-, 7- or 14-days post-inoculation (dpi). Prior to snap-freezing, extraradical fungal mycelium was removed by washing the roots in ice-cold water.

### RNA extraction, cDNA synthesis and qRT-PCR

Plant root tissue or fungal mycelium was ground to a fine powder in liquid nitrogen using a mortar and pestle, then transferred to 2 mL microcentrifuge tubes. Total RNA was extracted by adding 1 mL TRIzol® (ThermoFisher, # 15596018), followed by vigorous vortexing until the tissue was fully suspended. Subsequently, 200 µL chloroform were added, and the samples were vortexed for 20 sec. Phase separation was achieved by centrifugation at 13,000 rpm for 30 minutes at 4 °C. The aqueous phase was transferred to a fresh 1.5 mL tube and mixed with 500 µL of 100% isopropanol. RNA was precipitated overnight at -20 °C. The following day,

RNA was pelleted by centrifugation for 30 min at 13,000 rpm and 4 °C. The pellet was washed three times with 1mL 75 % (v/v) ethanol. After complete removal of ethanol, the pellet was dissolved in 30 µL RNAse-free water and incubated at 65 °C for 5 min. To remove co-precipitated genomic DNA, 10 µL DNase mix (4 µL 10× DNase buffer, 1 µL DNase I [1 U µL^-1^, Thermo Fischer, # EN0521], 5 µL MQ-H₂O) was added, and samples were incubated for 30 minutes at 37 °C. Following DNA digestion, 20 µL of 7.5 M NH^4+^-acetate and 80 µL of 100 % isopropanol were added to each tube. RNA was again precipitated overnight at -20 °C, pelleted by centrifugation, washed with 70 % (v/v) ethanol, air-dried and resuspended in 30 µL RNAse-free water by incubation at 65 °C for 5 min. RNA concentration and purity were assessed using a Nanodrop spectrophotometer and up to 2 µg of RNA were used for cDNA synthesis with the First Strand cDNA Synthesis Kit (ThermoFisher, #K1612) following the manufacturer’s instructions. qRT-PCR was performed using the GOTaq® qPCR Master Mix (Promega, #A6001) with gene-specific primers listed in Supplementary Table 1.

### Phylogenetic Analysis of ROCX sequences and HMM annotation

The 114 fungal ROC sequences previously reported in Marian et al. (2022) and the *Si*ROCX sequence (PIIN_00682) were used for phylogenetic analysis. In addition, an HMM profile search of *Si*ROCX against fungal ROCX sequences was done, and the top 8 hits were included. This resulted in 123 sequences in total. The sequences were aligned using MAAFT v7.0 (Katoh et al., 2002). Uninformative sites were trimmed using Clipkit v2.6.1 (Steenwyk et al., 2020) on the Linux Terminal. The processed sequences were used as an input for IQ-TREE v2.0.7 (Minh et al., 2020) on the terminal. First, model inference without tree reconstruction was carried out using the -MF argument to select the best evolutionary model. Tree reconstruction was done using the LG+F+R7 model. Branch support using Ultrafast bootstrapping was calculated with the argument -B 1000, and 31 threads were used with -T 31. The sequences were annotated for the presence of the ROCX HMM motif. For this purpose, ROCX sequences from Agaricomycetes were used to build an HMM profile (Marian et al., 2022). 63 sequences were used an input for the hmmbuild function from the hmmer package v3.3.2 in the Linux terminal (Eddy, 2011). The resulting HMM profile was aligned to the fungal sequences used for the tree using hmmsearch from the same package, and the sequences with a best domain score > 500 were considered positive for the ROCX HMM motif.

## Data availability

All data supporting the findings of this study are available within the article and supplementary material.

## Acknowledgements

We acknowledge the work (proposal: 10.46936/10.25585/60001292) conducted by the US Department of Energy Joint Genome Institute (https://ror.org/04xm1d337), a DOE Office of Science User Facility, is supported by the Office of Science of the US Department of Energy operated under contract no.: DE-AC02-05CH11231.

We thank the CECAD Proteomics Core Facility for analysis and support during proteomics experiment. This work was supported by the large instrument grant INST 216/1163-1 FUGG by the German Research Foundation (DFG Großgeräteantrag).

AZ, VR, MP and LA acknowledge support from the Cluster of Excellence on Plant Sciences (CEPLAS), funded by the Deutsche Forschungsgemeinschaft (DFG, German Research Foundation) under Germany’s Excellence Strategy – EXC 2048/1, Project ID 390686111. FD and AZ acknowledge support from the research unit FOR5682 funded by the Deutsche Forschungsgemeinschaft (DFG, German Research Foundation) under the Project ID 520490591.

## Author contributions

AZ, MB and LA conceived and designed the research. LA, MB and AZ wrote the manuscript. LA, FD, VR and MB performed the experiments. LA, FD, VR, MP and MB analyzed the data. LA conducted the bioinformatic analysis of co-regulated genes. AZ provided funding for the experiments. All authors contributed to manuscript editing and approved the final version.

## Competing interests

The authors declare not competing interests.

## Supplementary Figures

**Supplementary Fig. 1.**
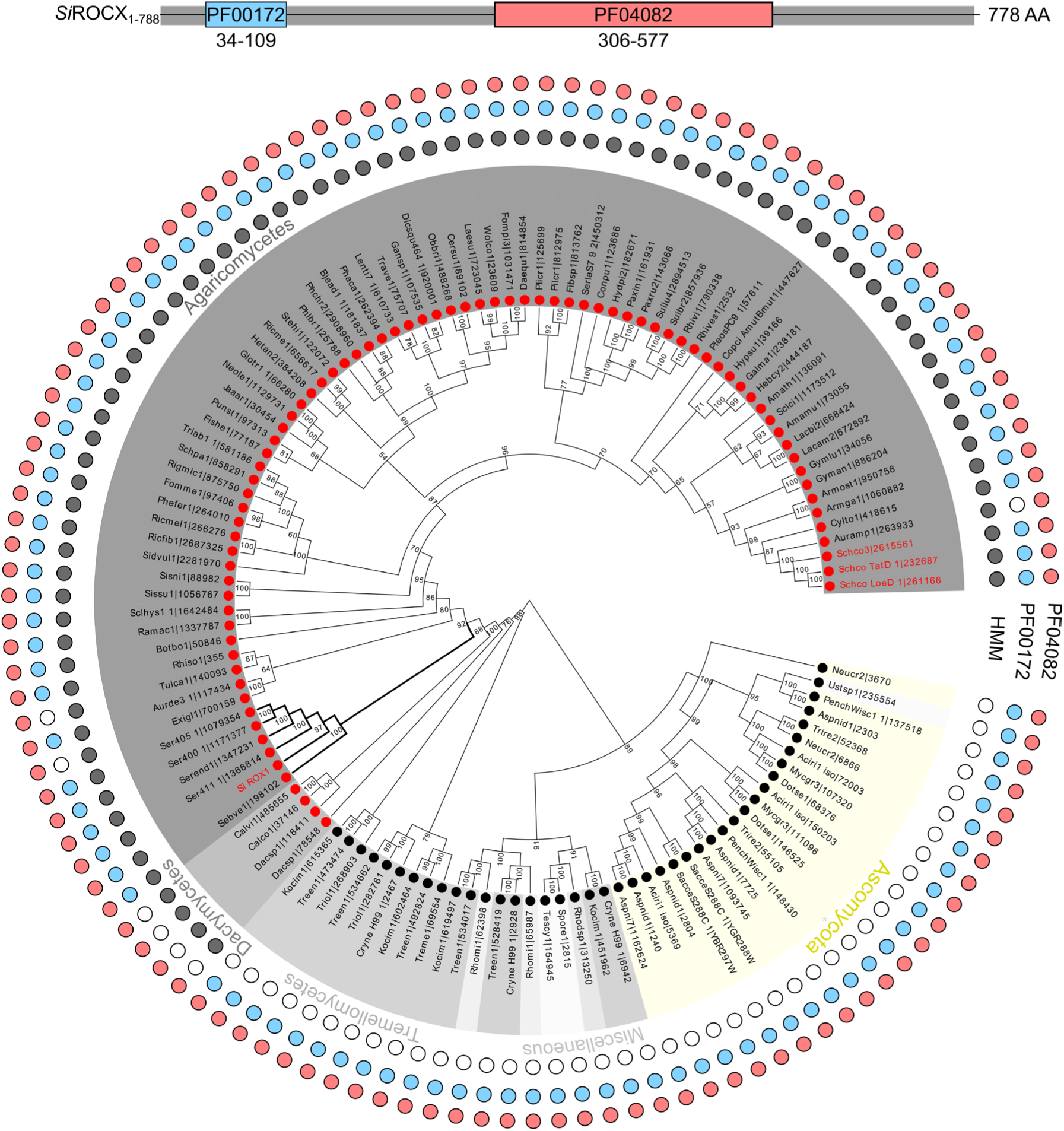
Domain architecture and phylogeny of *Si*ROCX and related fungal Zn₂Cys₆ transcription factors. The domain architecture is shown for the manually curated full-length *Si*ROCX protein (PIIN_00682; 788 aa). Conserved domains were identified using Pfam and include the Zn₂Cys₆ DNA-binding domain (PF00172) and the fungal-specific transcription factor domain (PF04082). The phylogeny shows the relationships among 114 fungal sequences reported by Marian et al. (2022), together with *Si*ROCX and additional closely related sequences identified by HMM search. Background colors indicate taxonomic affiliation (yellow = Ascomycota; grey = Basidiomycota). Different shades of grey denote major Basidiomycota classes (including *Ustilaginomycetes*, *Cystobasidiomycetes*, and *Exobasidiomycetes*). Colored rings indicate presence of conserved domains and HMM signatures: ROCX-specific HMM profile (black), Pfam domain PF00172 (Zn₂Cys₆ DNA-binding domain; blue), and PF04082 (fungal-specific transcription factor domain; red). ROCX sequences from *Serendipita indica* and *Schizophyllum commune* are highlighted in red. Node labels indicate bootstrap support values, and branches with bootstrap support >95 are considered well supported.

**Supplementary Fig. 2.**
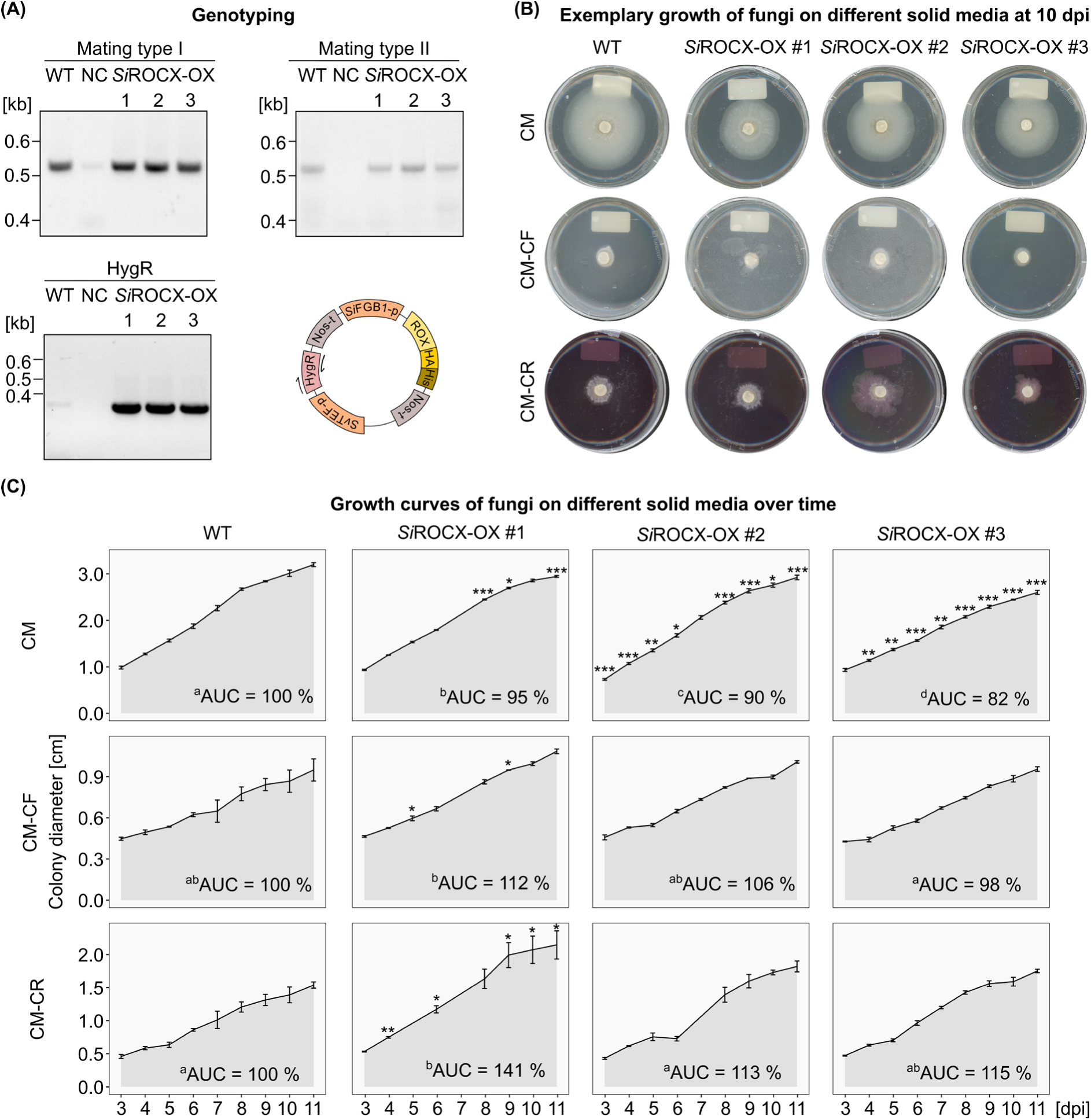
Characterization of *Si*ROCX overexpression strains. **A)** PCR-based genotyping of wild-type (WT) *Serendipita indica* and three independent *Si*ROCX overexpression transformants confirming mating type and integration of the transgene. A schematic representation of the transformation construct used for *Si*ROCX overexpression is shown. **B)** Growth of fungal strains on CM agar or CM agar supplemented with Calcofluor White (CF) or Congo Red (CR) at 10 dpi. Representative colonies are shown. **C)** Colony growth quantified from 3–11 dpi for strains grown on CM agar or CM agar supplemented with CF or CR. Growth curves represent mean colony diameter (n = 3). Area under the curve (AUC) values were calculated for each genotype and compared using one-way ANOVA followed by Tukey’s post hoc test (p < 0.05). Different letters indicate significant differences between genotypes. For individual time points, significant differences relative to WT are indicated by asterisks (* p < 0.05; ** p < 0.01; *** p < 0.001).

## Notes

### Competing Interest Statement

The authors have declared no competing interest.

